# HGTs are not SPRs: In the presence of ghost lineages, series of Horizontal Gene Transfers do not result in series of Subtree Pruning and Regrafting

**DOI:** 10.1101/2024.11.22.624805

**Authors:** Eric Tannier, Théo Tricou, Syrine Benali, Damien M. de Vienne

**Author notes:** Corresponding author: DAMIEN M. DE VIENNE, Université Lyon 1, CNRS, Laboratoire de Biométrie et Biologie Évolutive, Bâtiment Mendel 43 boulevard du 11 Novembre 1918, 69622 VILLEURBANNE CEDEX.

## Abstract

When a gene is horizontally transferred (HGT), under the “replacement” model where the transferred gene replaces its homolog in the recipient genome, the corresponding gene phylogeny departs from the species phylogeny by a Subtree Prune and Regraft (SPR) operation: the “recipient branch” is moved from its initial position to attach to the “donor branch”. Based on this observation, various methods have used SPRs to infer HGTs. We examine this apparent equivalence in the light of ghost lineages, *i*.*e*. related species absent from the phylogeny because they are extinct, unknown or have not been sampled. In this case an SPR is not directly interpretable by an HGT from the donor branch, because HGTs can have ghost lineages as donors. A possible and frequent interpretation – that we call “induced HGT” – is that the transferred gene leaves the sampled phylogeny for a ghost lineage at the donor branch, and is transferred back from a ghost lineage at the recipient branch. We show by simulations that this interpretation is misleading in a significant number of cases. For instance if the studied phylogeny represents 1% of all the species susceptible to exchange genetic material with the 100 sampled species, and 11 transfers occurred, then SPRs do not correspond to induced HGTs in around 50% of the cases. This leaves the question of a coherent interpretation of SPR in the presence of ghosts open, and applies to a certain extent to other phylogenetic simulation or inference methods of HGT, like reconciliation, or phylogenetic networks.

## Introduction

Horizontal (or lateral) transfers depict the transmission of genetic material from one organism to another that is not its direct descendant, as opposed to the vertical transmission. This mode of transmission is particularly common in bacteria and archaea, where it is an essential evolutionary process that contributes to the adaptation of lineages to their environment (Arnold et al. 2022; Gophna and Altman-Price 2022). In eukaryotes, it is being increasingly recognized as an important process as well (Keeling 2024), though its significance largely remains to be explored.

The horizontal transfer of a gene (HGT) has a theoretical effect on phylogenies, which is commonly exploited for its detection (Ravenhall et al. 2015): if we represent the evolutionary history of genomes by a species phylogeny and the evolutionary history of a considered gene by a gene tree, an HGT can introduce a difference between the two trees. Indeed, consider a simple model where genes do not undergo duplications, losses or other events that can depart their evolutionary history from that of the species, and where an HGT consists of the replacement of a gene by its homolog in another lineage (it is a replacing transfer, Choi et al. 2012). In this model an HGT results in a Subtree Prune and Regraft (SPR) operation (Allen and Steel 2001): the gene tree topology resulting from the HGT can be obtained from the species tree topology by cutting (pruning) the recipient branch of the transfer and reattaching it (regrafting) at the location of the donor branch.

This correspondence between HGT and SPR is the basis of several HGT detection and simulation methods using transformation of trees by SPR operations (Ravenhall et al. 2015; Chan et al. 2017). Various algorithms and methods have been designed to determine sequences of SPR operations (or “edit paths”) that transform species trees into gene trees (Hallett and Lagergren 2001; Nakhleh et al. 2005; Suchard 2005; Beiko and Hamilton 2006; Hill et al. 2010; Avni and Snir 2020), such edit paths being interpreted in terms of horizontal transfers (e.g. Beiko 2005; Whidden et al. 2014). Tools have also been developed to simulate gene trees by performing SPRs in the species tree following HGTs (e.g., Horiike et al. 2011; Mallo et al. 2016).

We examine this correspondence between HGT and SPR in the light of an important parameter of the study of life evolution: the fact that we don’t have access to most species that live or have lived on earth. Indeed, the vast majority of species are either extinct (Raup 1991) or extant but unknown (Mora et al. 2011; Locey and Lennon 2016), and even for known species, only a tiny fraction is susceptible to be included in any analysis carried out. Phylogenetically speaking, this translates to phylogenetic trees often representing the evolutionary history of a small subset of all the species that descended from their root. Absent species or clades — referred to as ghost lineages — were shown to be of particular importance for the study of horizontal genetic flows (Durvasula and Sankararaman 2020; Ottenburghs 2020; Tricou et al. 2022a, 2022b) because given their overwhelming abondance, it is likely that they are involved, as donors or recipients, in some (or most) of the detected HGTs (Macleod et al. 2005; Galtier and Daubin 2008; Fournier et al. 2009; Szöllősi et al. 2013). The significant influence of ghost lineages on other fields of evolutionary biology such as historical biogeography (Faurby et al. 2024) or ancestral state reconstruction (Finarelli and Flynn 2006) was also acknowledged. The present study is a new contribution to this growing body of research.

### The importance of ghosts when interpreting SPRs as HGTs - Illustration

We propose to investigate the impact of ghost lineages on the robustness of the interpretation of SPRs as HGTs. This requires to differentiate “full trees”, *i*.*e*. species tree and gene tree including ghosts, and “sampled trees”, *i*.*e*. the species and gene trees from which ghost lineages have been pruned. In an evolutionary study one can only have access to sampled trees ; full trees are theoretical objects with many more species (extinct and extant).

As noted above, in full trees, HGTs are equivalent to SPRs. The pruned branch of the SPR is the recipient branch of the HGT, and the branch where it is regrafted is the donor branch of the HGT. In sampled trees this equivalence does not immediately hold because involved branches might be unsampled. First, let’s consider a scenario with only one HGT. If its recipient branch is a ghost then the HGT is simply not visible in the sampled trees. If the donor branch is a ghost (Figure 1A), then the sampled gene tree is obtained from the sampled species tree by an SPR, which pruned branch is still the recipient branch of the HGT, and which destination branch of the regrafting is the most recent ancestor, among sampled branches, of the (ghost) donor of the HGT. Indeed, regrafting on a ghost branch is equivalent, in the sampled tree, to regrafting to the ancestor of this ghost branch.

**Figure 1.**
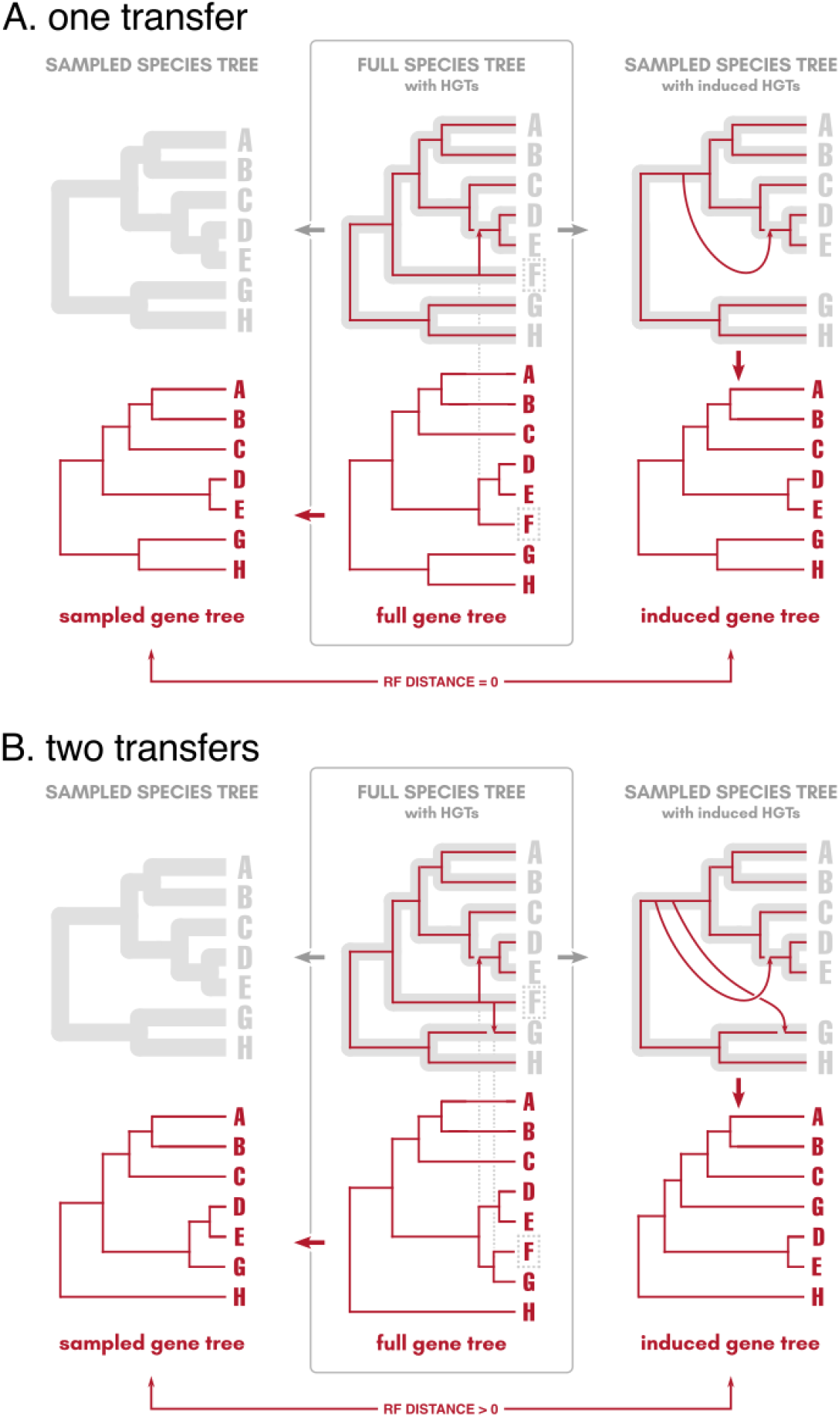
Illustration of the effect of ghosts on the possible disconnection between HGTs and SPRs. Each panel (**A** and **B**) is organised the same way, the only difference is the number of HGTs represented. For each panel, from a full species tree and a set of HGTs (top of the central box), it is possible to obtain the full gene tree (bottom of the central box) by performing SPRs on the full species tree following the HGTs. From the full species tree and the full gene tree (central box), it is direct to obtain the sampled species tree and the sampled gene tree (left part) by pruning the ghost species (F in this example) in both trees. These sampled trees are the only ones that can be observed in biological studies. From the full species tree and the known HGTs, it is also possible to compute the “induced HGTs” (top right), *i*.*e*. the HGTs in the sampled tree that correspond to the HGTs in the full tree. “Induced HGTs” correspond to mapping HGTs into the sampled species tree by identifying what branches in the sampled species tree are the donors and recipient branches of the HGTs present in the full species tree (see text). They are used as the interpretation of inferred SPRs. The induced gene tree is the gene tree resulting from performing SPRs in the sampled species tree following the induced HGTs. When more than one transfer involves the same ghost donor branch (panel **B**), performing these SPR operations leads to a gene tree that is topologically different from the sampled gene tree (RF distance between the two trees is not null). This is not the case when a single transfer (panel **A**) exists.

This is how an SPR is usually interpreted in the presence of ghosts: it corresponds to a sequence of evolutionary events, in which a gene leaves the sampled phylogeny for a ghost lineage attached to the donor branch in the species tree (following a speciation or a transfer, called “branching out” by Duchemin et al. 2018), evolves along ghost lineages, and a descendant of this gene is transferred from a ghost lineage to the recipient branch. The donor and recipient branches identified by the SPR are the branches where the gene leaves and re-enters the sampled phylogeny, and can be defined as the donor and recipient branches of an “induced HGT”. This interpretation of a transfer in the sampled tree as an induced HGT, summarising a sequence of events involving ghost lineages, is present in several evolutionary studies and methods on transfers or introgressions (Moret et al. 2004; Doyon et al. 2010; Yu et al. 2011; Szöllősi et al. 2013; Jacox et al. 2016). As an illustration, an induced HGT is defined as follows in Figure 1A: the HGT from F to (DE) becomes the “induced HGT” from (ABCDE) to (DE), *i*.*e*. from the branch in the sampled species tree that was initially supporting the ghost lineage (F in this example).

This interpretation of SPR as induced HGT holds for scenarios with only one HGT, but is not generalisable. We show it with the toy examples of Figure 1 where we compare two gene trees. One is the “sampled gene tree” (Figure 1A and B, bottom left) obtained by applying SPRs in the full tree and then sampling the non ghost lineages. The other is the “induced gene tree” (Figure 1A and B, bottom right) obtained by sampling the non ghost lineages in the species tree and then applying SPRs according to the induced HGTs. Under the hypothesis that a SPR should be interpreted as an induced HGT, these two trees should be identical. It is the case when a single HGT occurred, because regrafting to a ghost lineage is equivalent, in the sampled tree, to regrafting to its ancestor branch, up to the most recent sampled ancestor (Figure 1A). However, the situation is different when two or more transfers emerge from the same ghost branch (Figure 1B).

Translating the two HGTs in the full species tree into “induced HGTs” in the sampled species tree produces the two transfers highlighted in the top right of Figure 1B: the transfers F to (DE) and F to G have (ABCDE) to (DE) and (ABCDE) to G as induced transfers, because the new donor branch of the transfer is the branch in the sampled tree on which the ghost species F is attached in the full species tree. We observe that performing SPRs on the species tree following these induced transfers produces a gene tree (the induced gene tree in Figure 1B, bottom right) that is topologically different from the sampled gene tree (Figure 1B, bottom left). Thus, the interpretation of SPRs as induced HGTs does not hold as soon as there are two transfers from the same ghost clade.

### The importance of ghosts when interpreting SPRs as HGTs - Quantification

We use simulations (code and data available at https://github.com/damiendevienne/hgts-are-not-sprs) to explore and quantify the possible effect of this disconnection between HGTs and SPRs due to ghosts. This approaches the possible error rate of a method using SPRs to detect or simulate HGTs. We applied HGTs in full simulated species trees, computed the sampled gene tree and induced gene tree, and the proportion of cases where the trees are different is used as an estimation of the error rate of interpreting SPRs as induced HGTs. We applied this procedure instead of a more direct one, inferring SPRs on sampled trees and computing the donor error rate, to avoid the variability due to inference uncertainties and errors, as well as to save computing time. We repeated (100 replicates) the following operations for different numbers of transfers (t=2, 5, 8, 11, 14, 17 or 20) and different numbers of species sampled (n=50, 100, 500 or 1000, corresponding to 0.5%, 1%, 5% or 10% of the total number of species):

- Simulate a tree with 10 000 tips (full species tree T)
- Randomly sample *n* tips as being non-ghosts to produce the sampled species tree S
- Randomly choose donor and recipient branches (contemporaneous in the full tree) of *t* HGTs, ensuring that each transfer has an extant branch as recipient
- Compute the “sampled” gene tree, by
  ∘ Performing SPRs on T following the HGTs
  ∘ Sampling the *n* tips on the obtained tree
- Compute the “induced” gene tree, by
  ∘ Computing the induced HGTs based on *t, n* and N
  ∘ Performing SPRs on S following the induced HGTs
- Compute the topological distance (RF distance, Robinson and Foulds 1981) between the two gene trees

We observe (Figure 2) that the more ghosts there are, the more likely it is that the sampled and the induced gene trees will topologically disagree. It also appears that the proportion of cases where there is an incongruence increases with the number of HGTs, which was expected. If 10% of the species are sampled (90% of the species are ghosts) and there are 11 transfers involving an extant branch as recipient, around 7% of the sampled and induced gene trees will disagree. This percentage increases to around 50% if only 1% of the species are sampled (99% of the species are ghosts) and to 70% if 0.5% are sampled (Figure 2).

**Figure 2.**
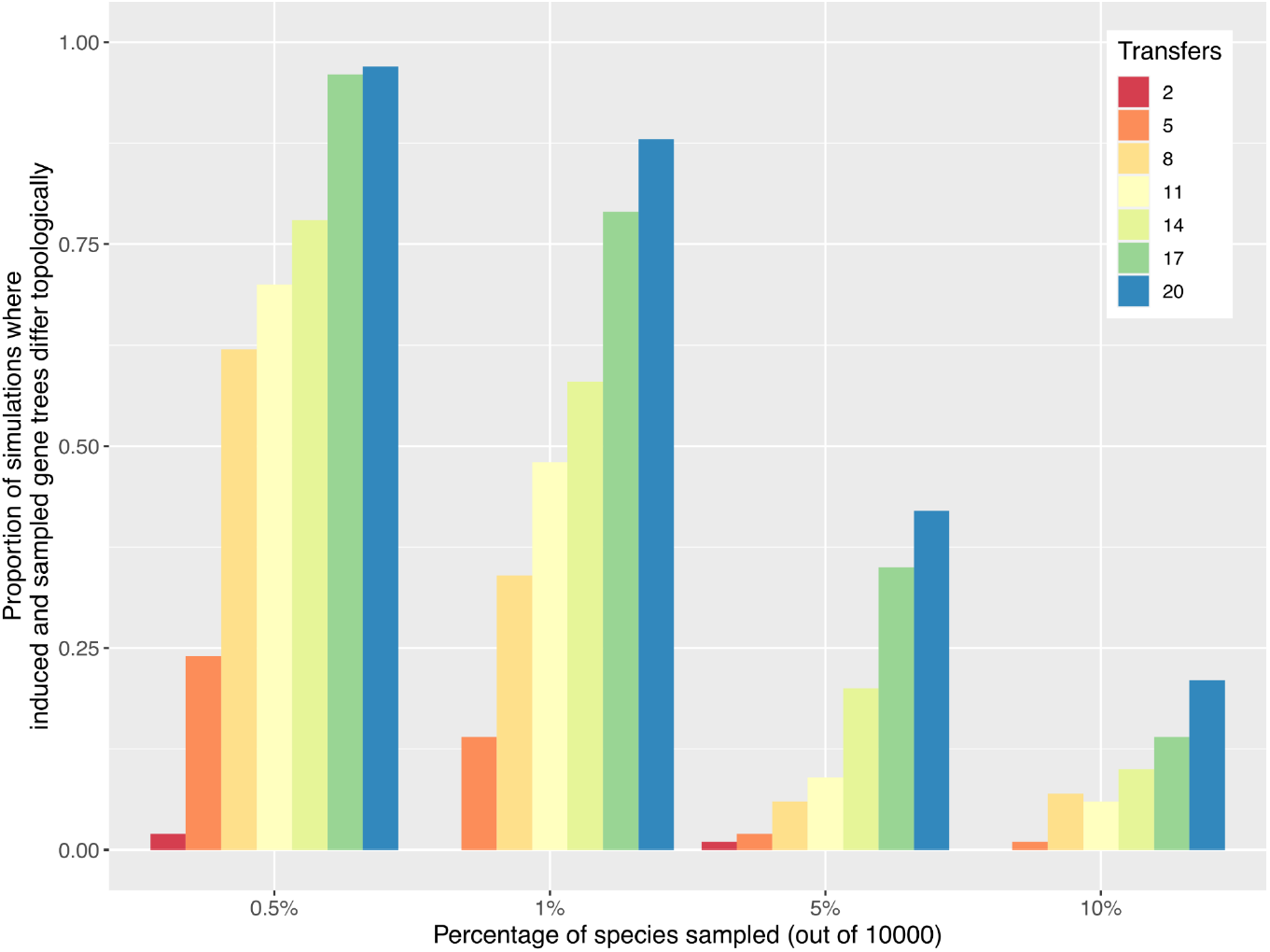
Effect of the amount of ghost lineages (x-axis) and of the number of horizontal gene transfers (HGTs) on the proportion of cases where the sampled gene trees and the induced gene trees (see text) are topologically different. This proportion approximates the possible error rate of a method using SPRs to detect or simulate HGTs.

Relating this to methods that use SPRs to infer HGTs, this means that if the species tree on which the method is used represents 1% of the total number of species that really populate this tree, and if 11 HGTs really occurred, trying to interpret the SPRs in terms of HGTs will be misleading 50% of the time because SPRs on the tree are not expected to produce the observed gene tree.

Looking at the RF distances between the true sampled gene tree and the induced gene tree in each case (Supplementary Figure 1), we observe that the trees not only often disagree but also sometimes quite substantially. For example, with 5 transfers on trees representing 1% of the total number of species (*i*.*e*. 100 tips in each tree), the RF distance reached up to 46.

## Conclusions

Our results show that in the presence of ghosts, series of SPRs transforming the extant species tree into the extant gene tree cannot be interpreted in terms of the induced HGT. As a consequence, a method transforming phylogenetic trees by the means of SPR cannot detect the donor branch if the species tree is not exhaustive (contains no ghosts). This calls for alternative interpretation of SPR edit paths in terms of HGTs.

To obtain our results, we defined the concept of “induced HGTs”, i.e. horizontal transfers in a given sampled species tree corresponding to the horizontal transfers occurring in the full species tree. Note that this concept, which should be useful for future research on the impact of ghosts for the study of gene flow, is present in the HGT inference literature. There, it is often termed “transfers to the future” and depicts the combination of a speciation to an unsampled lineage followed by a transfer back to a sampled one (see Menet et al. 2022 for a recent review).

In this study, we only quantify the problem of interpreting SPRs as induced transfers, but we do not provide a solution to the problem. Further work is therefore needed, which in turn could help in the identification and phylogenetic placement of ghost lineages in sampled species trees. We anticipate that the solution might be found more easily with recent reconciliation methods, such as DTL methods (which infer Duplications, Transfers and Losses from the comparison between gene and species trees) (Szöllősi et al. 2015; Jacox et al. 2016) that theoretically allow for diversification of genes outside the given species tree. However few software appear to have implemented this possibility, so inference methods in their current form might be subject to the same error risks.

## Supplementary material

**Supplementary Figure 1.**
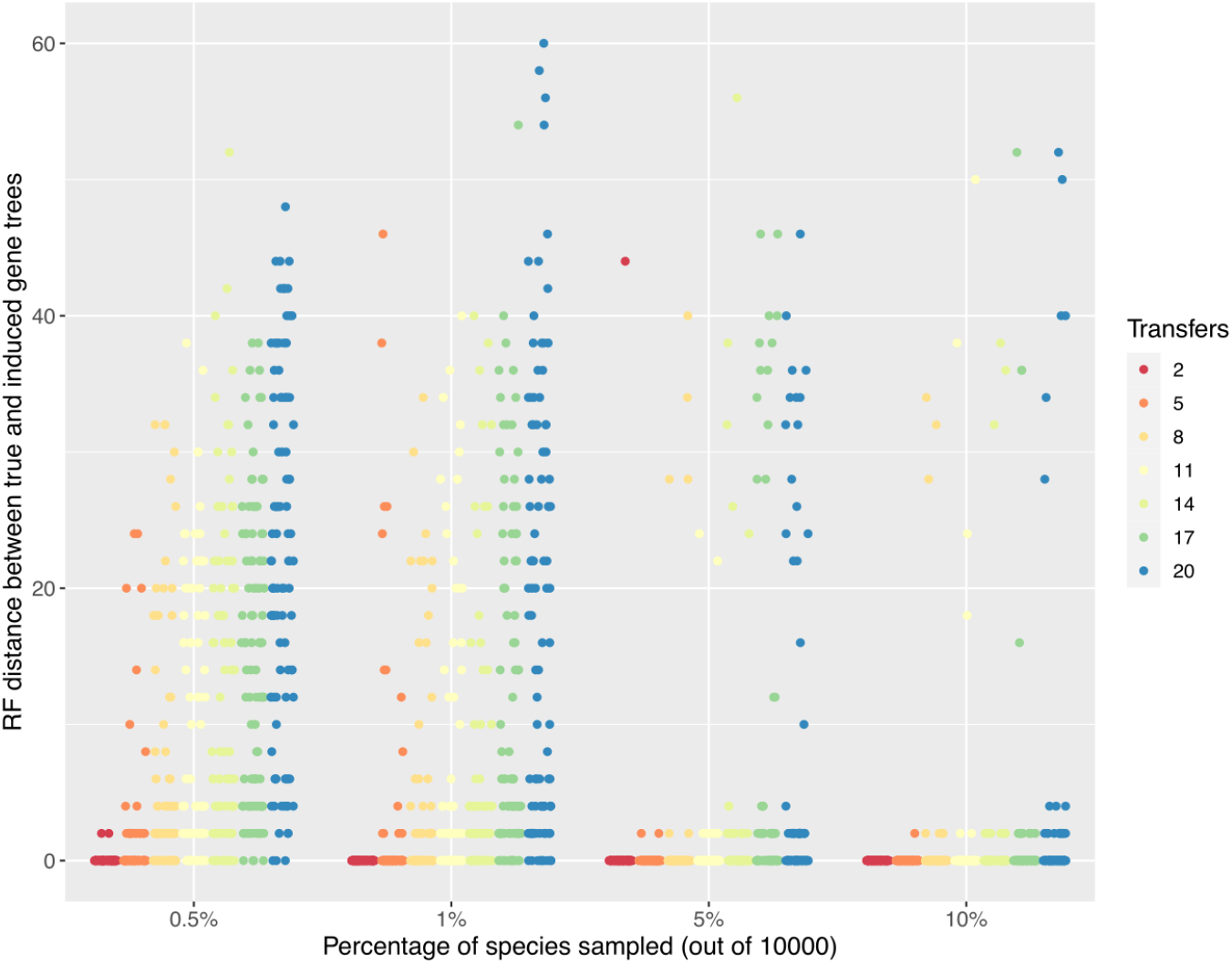
Effect of the amount of ghost lineages (x-axis) and of the number of horizontal gene transfers (HGTs) on the topological distance (RF distance, y-axis) between the sampled gene trees and the induced gene trees. Dots in each color represent replicates (N=100).

## References

Allen B.L., Steel M. 2001. Subtree Transfer Operations and Their Induced Metrics on Evolutionary Trees. Annals of Combinatorics. 5:1–15.

Arnold B.J., Huang I.-T., Hanage W.P. 2022. Horizontal gene transfer and adaptive evolution in bacteria. Nature Reviews Microbiology. 20:206–218.

Avni E., Snir S. 2020. A New Phylogenomic Approach For Quantifying Horizontal Gene Transfer Trends in Prokaryotes. Scientific Reports. 10:12425.

Beiko R.G. 2005. Highways of gene sharing in prokaryotes. Proceedings of the National Academy of Sciences. 102:14332–14337.

Beiko R.G., Hamilton N. 2006. Phylogenetic identification of lateral genetic transfer events. BMC Evolutionary Biology. 6:15.

Chan C.X., Beiko R.G., Ragan M.A. 2017. Scaling Up the Phylogenetic Detection of Lateral Gene Transfer Events. In: Keith J.M., editor. Bioinformatics: Volume I: Data, Sequence Analysis, and Evolution. New York, NY: Springer. p. 421–432.

Choi S.C., Rasmussen M.D., Hubisz M.J., Gronau I., Stanhope M.J., Siepel A. 2012. Replacing and Additive Horizontal Gene Transfer in Streptococcus. Molecular Biology and Evolution. 29:3309–3320.

Doyon J.-P., Scornavacca C., Gorbunov K.Y., Szöll\Hosi G.J., Ranwez V., Berry V. 2010. An efficient algorithm for gene/species trees parsimonious reconciliation with losses, duplications and transfers. Comparative Genomics: International Workshop, RECOMB-CG 2010, Ottawa, Canada, October 9-11, 2010. Proceedings 8.:93–108.

Duchemin W., Gence G., Arigon Chifolleau A.-M., Arvestad L., Bansal M.S., Berry V., Boussau B., Chevenet F., Comte N., Davín A.A., Dessimoz C., Dylus D., Hasic D., Mallo D., Planel R., Posada D., Scornavacca C., Szöllősi G., Zhang L., Tannier É., Daubin V. 2018. RecPhyloXML: a format for reconciled gene trees. Bioinformatics. 34:3646–3652.

Durvasula A., Sankararaman S. 2020. Recovering signals of ghost archaic introgression in African populations. Science Advances. 6:eaax5097.

Faurby S., Silvestro D., Werdelin L., Antonelli A. 2024. Reliable biogeography requires fossils: insights from a new species-level phylogeny of extinct and living carnivores. Proceedings of the Royal Society B: Biological Sciences. 291:20240473.

Finarelli J.A., Flynn J.J. 2006. Ancestral State Reconstruction of Body Size in the Caniformia (Carnivora, Mammalia): The Effects of Incorporating Data from the Fossil Record. Systematic Biology. 55:301–313.

Fournier G.P., Huang J., Gogarten J.P. 2009. Horizontal gene transfer from extinct and extant lineages: biological innovation and the coral of life. Philosophical Transactions of the Royal Society B: Biological Sciences. 364:2229–2239.

Galtier N., Daubin V. 2008. Dealing with incongruence in phylogenomic analyses. Philos Trans R Soc Lond, B, Biol Sci. 363:4023–9.

Gophna U., Altman-Price N. 2022. Horizontal Gene Transfer in Archaea—From Mechanisms to Genome Evolution. Annual Review of Microbiology. 76:481–502.

Hallett M.T., Lagergren J. 2001. Efficient algorithms for lateral gene transfer problems. Proceedings of the fifth annual international conference on Computational biology.:149–156.

Hill T., Nordstrom K.J., Thollesson M., Safstrom T.M., Vernersson A.K., Fredriksson R., Schioth H.B. 2010. SPRIT: Identifying horizontal gene transfer in rooted phylogenetic trees. BMC Evol Biol. 10:42.

Horiike T., Miyata D., Tateno Y., Minai R. 2011. HGT-Gen: a tool for generating a phylogenetic tree with horizontal gene transfer. Bioinformation. 7:211–213.

Jacox E., Chauve C., Szöllősi G.J., Ponty Y., Scornavacca C. 2016. ecceTERA: comprehensive gene tree-species tree reconciliation using parsimony. Bioinformatics. 32:2056–2058.

Keeling P.J. 2024. Horizontal gene transfer in eukaryotes: aligning theory with data. Nature Reviews Genetics. 25:416–430.

Locey K.J., Lennon J.T. 2016. Scaling laws predict global microbial diversity. Proceedings of the National Academy of Sciences. 113:5970–5975.

Macleod D., Charlebois R.L., Doolittle F., Bapteste E. 2005. Deduction of probable events of lateral gene transfer through comparison of phylogenetic trees by recursive consolidation and rearrangement. BMC Evol Biol. 5:27.

Mallo D., De Oliveira Martins L., Posada D. 2016. SimPhy : Phylogenomic Simulation of Gene, Locus, and Species Trees. Systematic Biology. 65:334–344.

Menet H., Daubin V., Tannier E. 2022. Phylogenetic reconciliation. PLOS Computational Biology. 18:e1010621.

Mora C., Tittensor D.P., Adl S., Simpson A.G.B., Worm B. 2011. How Many Species Are There on Earth and in the Ocean? PLoS Biology. 9:e1001127.

Moret B.M.E., Nakhleh L., Warnow T., Linder C.R., Tholse A., Padolina A., Sun J., Timme R. 2004. Phylogenetic networks: modeling, reconstructibility, and accuracy. IEEE/ACM Transactions on Computational Biology and Bioinformatics. 1:13–23.

Nakhleh L., Ruths D., Wang L.-S. 2005. RIATA-HGT: A Fast and Accurate Heuristic for Reconstructing Horizontal Gene Transfer. Computing and Combinatorics.:84–93.

Ottenburghs J. 2020. Ghost Introgression: Spooky Gene Flow in the Distant Past. BioEssays. 42:2000012.

Raup D.M. 1991. Extinction: bad genes or bad luck? New York: W.W. Norton.

Ravenhall M., Škunca N., Lassalle F., Dessimoz C. 2015. Inferring horizontal gene transfer. PLoS computational biology. 11:e1004095.

Robinson D.F., Foulds L.R. 1981. Comparison of phylogenetic trees. Mathematical Biosciences. 53:131–147.

Suchard M.A. 2005. Stochastic Models for Horizontal Gene Transfer: Taking a Random Walk Through Tree Space. Genetics. 170:419–431.

Szöllősi G.J., Davín A.A., Tannier E., Daubin V., Boussau B. 2015. Genome-scale phylogenetic analysis finds extensive gene transfer among fungi. Philosophical Transactions of the Royal Society B: Biological Sciences. 370:20140335.

Szöllősi G.J., Tannier E., Lartillot N., Daubin V. 2013. Lateral Gene Transfer from the Dead. Systematic Biology. 62:386–397.

Tricou T., Tannier E., de Vienne D.M. 2022a. Ghost lineages highly influence the interpretation of introgression tests. Systematic Biology. 71:1147–1158.

Tricou T., Tannier E., de Vienne D.M. 2022b. Ghost lineages can invalidate or even reverse findings regarding gene flow. PLOS Biology. 20:1–16.

Whidden C., Zeh N., Beiko R.G. 2014. Supertrees Based on the Subtree Prune-and-Regraft Distance. Systematic Biology. 63:566–581.

Yu Y., Than C., Degnan J.H., Nakhleh L. 2011. Coalescent Histories on Phylogenetic Networks and Detection of Hybridization Despite Incomplete Lineage Sorting. Systematic Biology. 60:138–149.

